# Epilepsy-causing *KCNT1* variants increase K_Na_1.1 channel activity by disrupting the activation gate

**DOI:** 10.1101/2021.09.16.460601

**Authors:** Bethan A. Cole, Nadia Pilati, Jonathan D. Lippiat

## Abstract

Gain-of-function pathogenic missense *KCNT1* variants are associated with several developmental and epileptic encephalopathies (DEE). With few exceptions, patients are heterozygous and there is a paucity of mechanistic information about how pathogenic variants increase K_Na_1.1 channel activity and the behaviour of heterotetrameric channels comprising both wild-type (WT) and variant subunits. To better understand these, we selected a range of variants across the DEE spectrum, involving mutations in different protein domains and studied their functional properties. Whole-cell electrophysiology was used to characterise homomeric and heteromeric K_Na_1.1 channel assemblies carrying DEE-causing variants in the presence and absence of 10 mM intracellular sodium. Voltage-dependent activation of homomeric variant K_Na_1.1 assemblies were more hyperpolarised than WT K_Na_1.1 and, unlike WT K_Na_1.1, exhibited voltage-dependent activation in the absence of intracellular sodium. Heteromeric channels formed by co-expression of WT and variant K_Na_1.1 had activation kinetics intermediate of homomeric WT and variant K_Na_1.1 channels, with residual sodium-independent activity. In general, WT and variant K_Na_1.1 activation followed a single exponential, with time constants unaffected by voltage or sodium. Mutating the threonine in the K_Na_1.1 selectivity filter disrupted voltage-dependent activation, but sodium-dependence remained intact. Our findings suggest that K_Na_1.1 gating involves a sodium-dependent activation gate that modulates a voltage-dependent selectivity filter gate. Collectively, all DEE-associated K_Na_1.1 mutations lowered the energetic barrier for sodium-dependent activation, but some also had direct effects on selectivity filter gating. Destabilisation of the inactivated unliganded channel conformation can explain how DEE-causing amino acid substitutions in diverse regions of the channel structure all cause gain-of-function.

## Introduction

Epilepsy of infancy with migrating focal seizures (EIMFS) is a severe, pharmacoresistant developmental and epileptic encephalopathy (DEE). The disorder typically presents within the first six months of life, after which seizure frequency increases and is accompanied by other severe comorbidities such as developmental disorders ^1,2^ and delayed motor function ^3^. *De novo* gain-of-function (GOF) *KCNT1* pathogenic variants have been identified as the most frequent cause of EIMFS ^3,4^, and only one pathogenic variant thus far has been reported to cause a loss-of-function ^5^. *KCNT1* pathogenic variants, again GOF, have also been implicated in autosomal-dominant or sporadic sleep-related hypermotor epilepsy ((AD)SHE), a disorder characterised by motor seizures that occur during sleep ^6^. As with EIMFS, seizures are accompanied by psychiatric and intellectual disabilities. The mean age of onset for (AD)SHE is 6 years old; later than that for EIMFS ^1,6^. *KCNT1* pathogenic variants are also linked to other hyperexcitability disorders such as Ohtahara syndrome ^7^, Lennox-Gastaut syndrome ^8^, Status Dystonicus ^9^, West syndrome, leukoencephalopathies and Brugada syndrome ^2^. Pathogenic cardiac effects and ‘collateralopathies’ arising from *KCNT1* variants are becoming more widely reported ^10–12^.

*KCNT1* encodes the K_Na_1.1 subunit (Slack, or previously Slo2.2 or K_Ca_4.1), which forms a tetrameric potassium channel that is activated by intracellular Na^+^ and is widely distributed in the central nervous system. In the rat brain, immunoreactivity has been detected in the brainstem, olfactory bulb, frontal cortex, forebrain, thalamus, midbrain and cerebellum ^13–15^. K_Na_ channels, likely formed by K_Na_1.1 and closely related K_Na_1.2 (encoded by *KCNT2*) ^16,17^, have been implicated in the generation of the slow afterhyperpolarisation (AHP) following single action potentials ^18,19^. In principle neurons of the medial nucleus of the trapezoid body (MNTB), K_Na_ channels generate an AHP following bursts of action potential firing, regulating inter-burst timing and accuracy of firing ^20^. In addition to their role AHP, they are also important determinants of the resting membrane potential and intrinsic excitability in a number of cell types in the central nervous system (CNS) ^21,22^, and also outside of the CNS in arterial smooth muscle cells ^23^. Considering the role of K_Na_1.1 in neuronal excitability, how GOF pathogenic variants may lead to hyperexcitability is unclear. Several mechanisms have been proposed; for example, GOF of K_Na_1.1 channels in GABAergic inhibitory interneurons may increase hyperpolarisation and dampen their inhibitory effect on excitatory interneurons, leading to increased excitation ^24,25^. Another possibility is that variant K_Na_1.1 channels shorten the duration of action potentials and increase the AHP amplitude, increasing the rate of high frequency firing and resulting in hyperexcitability ^26^.

K_Na_1.1 is the largest known K^+^ channel subunit and has structural and functional similarities to K_Ca_1.1 (BK_Ca_, *KCNMA1*), the large conductance Ca^2+^-activated K^+^ channel, though sharing only ~7 % sequence homology ^13,16^. K_Na_1.1 subunits possess six transmembrane alpha helices (S1 to S6) and a pore-forming region between S5 and S6, containing the selectivity filter ^27,28^. The channel is weakly voltage-gated through a poorly understood mechanism. Unlike K_Ca_1.1 and voltage-gated K^+^ (K_v_) channels, K_Na_1.1 does not possess an excess of positively charged residues in its S4 transmembrane helices that confer voltage-sensitivity ^13,14^. Also unlike K_V_ channels, gating does not involve the opening of a transmembrane S6 helix bundle, but is likely to involve residues in or very close to the selectivity filter ^29,30^. This may resemble either the hydrophobic gating described for K_Ca_1.1 ^31^ or the selectivity filter gating mechanism identified in two pore domain K^+^ (K2P) channels ^32^. Analysis of the arrangements of the transmembrane helices between the inactive and active K_Na_1.1 conformations reveals that the only notable changes are in S6 and, to a lesser extent, S5 ^28^. This results in a change in the shape of the intracellular pore vestibule and could potentially couple the conformational changes induced by Na^+^ binding to the intracellular domains to channel opening. This mechanism of channel activation has been investigated using X-ray crystallography and molecular dynamics simulations of the prokaryotic MthK Ca^2+^-activated K^+^ channel ^33^, which shares structural features with K_Na_1.1. Here, binding of Ca^2+^ ions to the MthK RCK domains results in motion of the M1 and M2 helices (equivalent to S5 and S6 in K_Na_1.1) to open an activation gate, which is allosterically coupled to the selectivity filter gate, and resulting in K^+^ ion permeation ^33^. A similar mechanism could be at play in K_Na_1.1 and may be affected by structural changes caused by missense pathogenic variants.

The phenotypic spectrum of *KCNT1* pathogenic variants is broad and all of the reported DEE-causing pathogenic variants of *KCNT1*, some of which are recurring, are missense and generally result in increased channel activity ^34^. To date, substitutions of at least 60 of the amino acids in the K_Na_1.1 protein have been reported ^35^. There are locations within the channel structure where clusters of pathogenic variants have been identified; this is particularly evident around the RCK domains and NAD^+^ binding domain. However, some pathogenic variants are also found in the transmembrane regions close to the channel pore. Since the RCK1 and 2 domains and the S5-S6 pore-forming region are both thought to be involved in channel gating ^27,36,37^, it is likely that pathogenic variants interfere with gating in some way, such as by altering the Na^+^ sensitivity of the channel ^36–38^ and increasing probability of opening ^36^. Other studies report increased cooperative gating between variant channels ^3^, channels being in a constitutively phosphorylated-like state ^4^, and altered interactions with binding proteins such as Phactr1 ^39,40^. Importantly, DEE patients with *KCNT1* pathogenic variants are heterozygous, in general, carrying only one mutated allele. There is a lack of information on the behaviour of heteromeric channels comprising variant and wildtype (WT) subunits. Most functional studies of variant channels have been carried out by studying homomeric variant vs WT channel properties ^3–5,9,36,37,41,42^. Recently the implications of some heterozygous *KCNT1* variants ^38,43^ or *KCNT2* variant subunits co-expressed with *KCNT1* ^44^ on channel function have been assessed *in vitro*. These studies suggest that heteromeric variant+WT channels display ‘intermediate’ characteristics, with current amplitudes and activation parameters sitting between WT and variant homomeric channels.

DEEs resulting from pathogenic variants of *KCNT1* reported are mostly refractory ^45^ and could be treated using therapies that suppress overactive channel function. The antiarrhythmic drugs quinidine and bepridil have efficacy in inhibiting both WT and variant channels *in vitro* ^38,41,43,46^. Clofilium, another antiarrhythmic drug, also inhibits WT channels *in vitro* ^47^. Each of these drugs are non-selective and inhibit several cation channels ^38,46,47^ including hERG ^42,48^. This is the likely reason for the poor and inconsistent response in *KCNT1* patients when given quinidine therapy ^45^. Recently, more potent compounds have been identified, including some that have reduced potency in inhibiting hERG channels ^49–51^. However, to better understand how channel activity can be supressed pharmacologically, we require a better understanding of how the effects of all pathogenic variants converge on the K_Na_1.1 channel gating mechanism and how this persists in channel complexes containing both WT and variant subunits. Here, we selected seven different (AD)SHE or EIMFS-causing *KCNT1* variants that have been previously reported, have had different mechanisms proposed for GOF, and result in substitution of amino acids from across the K_Na_1.1 subunit structure (Fig. 1). We hypothesised that there would be common mechanisms underlying GOF in channels comprised of variant subunits alone and when co-expressed with WT. Since the WT channel is primarily dependent on intracellular Na^+^ for activation, and a number of variants reportedly alter the Na^+^ sensitivity of the channel ^36,38^, we have examined the effects of removing intracellular Na^+^ and the properties of unliganded channels. Our investigation led us to propose that all DEE-causing K_Na_1.1 variants disrupt the activation gate and destabilise its closed or inactive conformation, leading to enhanced activation upon stimulation.

**Figure 1:**
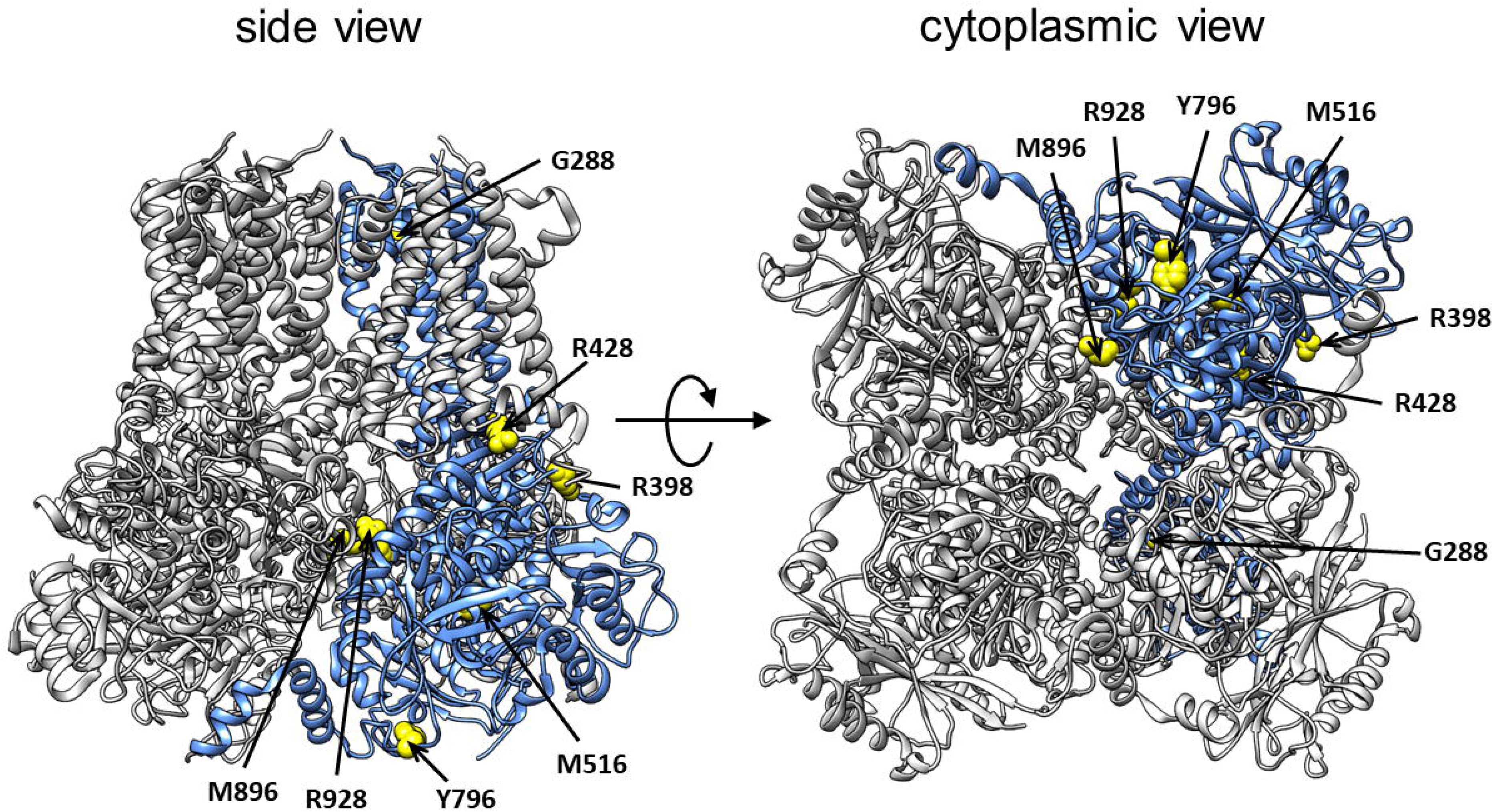
Location of amino acids mutated in this study in the structure of the chicken K_Na_1.1 channel in the active conformation (PDB: 5U70). One subunit of the K_Na_1.1 tetramer is coloured blue, with each of the amino acids mutated shown and coloured yellow. Views are from the side (left) with the transmembrane helices uppermost and the intracellular domains below, and the view from the cytoplasm (right). Figure prepared in UCSF Chimera.

## Materials and Methods

### Molecular biology and cell culture

A full-length human K_Na_1.1 cDNA clone in the pcDNA6-V5/His6 vector (Invitrogen) as used previously, was used in this study ^51^. Due to the high GC content of the *KCNT1* sequence, regions corresponding to the internal Bsu36I/BspEI, BspEi/SbfI, and SbfI/BsiWI restriction fragments were each amplified by PCR and cloned into the pJET1.2 cloning vector (CloneJET blunt cloning kit, Thermo Fisher Scientific, Loughborough, U.K.). Site directed mutagenesis was carried out on these constructs to introduce the (AD)SHE-causing Y796H, R928C, M896I and R398Q variants, EIMFS-causing R428Q, M516V and G288S variants, and selectivity filter mutant T314C using the NEB mutagenesis method (New England Biolabs, Hitchin, U.K.). Mutated sequences were verified by Sanger sequencing (Genewiz, Takely, U.K.) and subcloned into the pcDNA6-K_Na_1.1 construct at the corresponding restriction sites.

Chinese hamster ovary (CHO) cells were cultured in Dulbecco’s modified Eagle medium (DMEM) containing GlutaMax, and supplemented with 10 % Fetal Bovine Serum, 50 U/ml penicillin, and 0.5 mg/ml streptomycin (all from Thermo Fisher Scientific, Loughborough, U.K.) and incubated at 37 ºC in air containing 5% CO_2_. Cells were transiently transfected with WT or mutant K_Na_1.1 cDNA using Mirus Bio TransIT-X2 transfection reagent (Geneflow, Lichfield, U.K.). For homomeric expression 1.5 μg plasmid was used per 35 mm well. For heteromeric co-expression 0.75 μg of each construct was used. In all cases, a plasmid encoding Enhanced Green Fluorescent Protein (EGFP) (0.04 μg) was co-expressed as a marker of transfection. Cells were plated onto borosilicate glass cover slips and used for electrophysiological experiments 2-4 days post-transfection.

### Electrophysiology

All chemicals were obtained from Sigma-Aldrich (Gillingham, U.K.) unless otherwise stated. Macroscopic currents from CHO cells transiently transfected with mutant and WT K_Na_1.1 were obtained using the whole-cell configuration of the gigaseal patch clamp technique 2 - 4 days post-transfection, at room temperature (20 - 22ºC). Glass micropipettes of 1.5 - 2.5 MΩ resistance were pulled from thin-walled, filamented borosilicate glass capillaries (Harvard Apparatus Ltd, Edenbridge, U.K.), and fire-polished before filling with experimental solutions. The pipette solution contained, in mM, 100 K-gluconate, 30 KCl, 10 Na-gluconate, 29 glucose, 5 EGTA and 10 HEPES, pH 7.3 with KOH and the bath solution contained, in mM, 140 NaCl, 1 CaCl_2_, 5 KCl, 29 glucose, 10 HEPES and 1 MgCl_2_, pH 7.4 with NaOH. Na-gluconate in the intracellular solution was substituted with choline chloride for Na^+^-free experiments. Following establishment of a >2 GΩ seal, the cell membrane was ruptured to achieve the whole-cell configuration. Currents were recorded using a HEKA EPC 10 amplifier (HEKA electronic, Lambrecht, Germany) with 2.9 KHz low-pass filtering and 10 kHz digitisation. An Ag-AgCl electrode connected to the bath solution using a KCl-agar bridge as a reference. Data were collected using Patchmaster, with Fitmaster software (HEKA electronic, Lambrecht, Germany) used for offline analysis. To determine I-V relationships, the voltage protocol consisted of 400 ms steps from −100 to +80 mV in 10 mV increments, from a holding potential of −80 mV. Series resistance for whole cell recordings was <6 MΩ and compensated >65%.

### Data analysis

Data are presented as mean ± SEM of N number of cells. Statistical analysis was performed using SPSS (IBM Analytics, Portsmouth, U.K.), with the chosen tests indicated in figure legends; *p*<0.05 was considered significant. Representative whole-cell and excised inside-out current traces were plotted, and residual capacitance spikes removed in Origin Pro. Whole cell current-voltage relationships were divided by whole-cell capacitance to give current density (pA/pF). Reversal potentials were obtained by fitting the linear part of current-voltage relationships around the reversal potential using linear regression and determining the voltage at which current was 0 nA. Conductance (*G*) at each voltage (*V*_*m*_) was obtained by dividing current amplitudes by the driving force for K^+^ ions, calculated using the reversal potential (*V*_*rev*_) obtained in individual recordings:

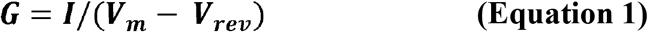

Conductance values were then plotted and fitted with a Boltzmann function:

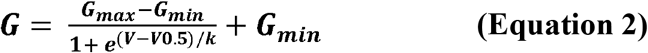

which gave values for half-maximal activation voltage (*V*_*0.5*_), *G*_*max*_, *G*_*min*_, and Slope factor (*k*). Data were normalised by dividing by *G*_*max*_ for each recording. With *k* = *RT/zF*, the elementary gating charge, *z*, was determined, and *ΔG*_*0*_, the zero-voltage free energy for channel opening, calculated from Δ*G*_*0*_ = *zFV*_*0.5*_. Currents were fit with a single exponential function at each voltage measured:

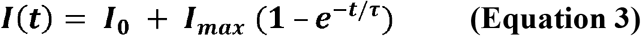

where *I*_*0*_ is the current offset, *I*_*max*_ is the steady-state current amplitude and r is the activation time constant.

## Results

### All homomeric *KCNT1* pathogenic variants lead to an increase in macroscopic current

Macroscopic currents through mutant K_Na_1.1 channels carrying EIMFS- and (AD)SHE-causing *KCNT1* variants were recorded using the whole cell configuration from CHO cells expressing either WT K_Na_1.1 or mutant channels. EIMFS-causing variants G288S, R428Q, and M516V, and (AD)SHE-causing variants Y796H, R928C, M896I, and R398Q were studied. Consistent with previous studies ^16,17,52^, WT K_Na_1.1 channels gave outward currents in response to a series of voltage steps from −100 mV to +80 mV from a holding potential of − 80 mV with 10 mM intracellular Na^+^ and the current consisted of an instantaneous component followed by a slower, time-dependent component (Figure 2A and 2B). Non-transfected CHO cells displayed very little outward current (1.49 ± 0.67 pA/pF, N=8, at +10 mV), with no time or voltage-dependence; implying negligible endogenous activity. In the absence of intracellular Na^+^, there was similarly no discernible WT K_Na_1.1 current across the same voltage range (2.93 ± 1.08 pA/pF, N=6, at +10 mV); confirming that intracellular Na^+^ is an absolute requirement for activation of the WT K_Na_1.1 channel under these conditions. With 10 mM intracellular Na^+^ and in agreement with previous reports ^3,4,6,9,37,38,41,43^, each of the mutant K_Na_1.1 channels examined resulted in large GOF (Figure 2A-C). EIMFS-causing mutants caused a 6-7-fold increase in mean peak current density at +10 mV (Figure 2B) compared to WT K_Na_1.1. (AD)SHE-causing mutants caused a 4-fold increase, except for Y796H, which resulted in a 6-fold increase (Figure 2B). From I-V relationships, mutations caused an increase in inward currents at membrane potentials negative to the reversal potential, as well as outward current and shifted activation towards more negative potentials (Figure 2C and Table 1). The zero-voltage activation energy change calculated for each mutant K_Na_1.1 yielded negative values, unlike the positive value obtained with WT (Table 1).

**Table 1:** Parameters derived from Boltzmann fit of WT, homomeric mutant, and co-expressed mutant+WT K_Na_1.1 channels in the absence and presence of intracellular 10 mM Na^+^, unless otherwise stated. Data are presented as mean ±SEM (N= 5-9 cells). V_0.5_ is the half-maximal activation voltage, \**p*<0.05, \*\**p*<0.005, \*\*\**p*<0.0005 compared to homomeric mutant with 10 mM [Na^+^]i (independent one-way ANOVA with Tukey’s post-hoc test). Elementary gating charge, *z,* was derived from the slope of the Boltzmann curve, *RT/zF*. ΔG_0_ is the zero-voltage free energy for channel opening (Δ***G***_0_ = ***zFV***_0.5_) and is expressed as kcal/mol. ΔΔG_0_ is the closed-state destabilisation energy, calculated as the difference in mean ΔG_0_ between WT and mutant K_Na_1.1 channels.

| Channel | V<sub>0.5</sub><br>(mV) |  | *z* | Δ*G*<sub>0</sub><br>(kcal/mol) | ΔΔ*G*<sub>0</sub><br>(kcal/mol) |
| --- | --- | --- | --- | --- | --- |
| WT | 21.77 ±4.35 |  | 1.01 ±0.13 | 0.49 ±0.10 | - |
| G288S | -20.31 ±8.77 |  | 0.66 ±0.09 | -0.36 ±0.18 | 0.85 |
| G288S + WT | 2.63 ±4.18 | * | 0.59 ±0.04 | 0.03 ±0.05 | 0.46 |
| G288S 0 mM [Na<sup>+</sup>]<sub>i</sub> | 12.89 ±5.70 | ** | 0.50 ±0.03 | 0.15 ±0.07 | 0.34 |
| Y796H | -25.60 ±2.27 |  | 0.92 ±0.08 | -0.53 ±0.05 | 1.02 |
| Y796H + WT | 28.72 ±4.76 | *** | 0.79 ±0.06 | 0.54 ±0.10 | -0.03 |
| Y796H 0 mM [Na<sup>+</sup>]<sub>i</sub> | -10.41 ±7.14 |  | 0.62 ±0.06 | -0.16 ±0.10 | 0.65 |
| R428Q | -0.22 ±6.19 |  | 0.73 ±0.03 | -0.02 ±0.11 | 0.51 |
| R428Q + WT | 20.20 ±2.97 | * | 0.69 ±0.05 | 0.31 ±0.03 | 0.18 |
| R428Q 0 mM [Na<sup>+</sup>]<sub>i</sub> | 44.12 ±5.33 | *** | 0.65 ±0.05 | 0.68 ±0.10 | -0.19 |
| M516V | -18.32 ±8.74 |  | 0.57 ±0.04 | -0.23 ±0.11 | 0.72 |
| M516V + WT | 21.47 ±6.57 | * | 0.66 ±0.05 | 0.35 ±0.10 | 0.14 |
| M516V 0 mM [Na<sup>+</sup>]<sub>i</sub> | 37.75 ±7.17 | ** | 0.65 ±0.04 | 0.57 ±0.11 | -0.08 |
| R398Q | -4.15 ±7.18 |  | 0.72 ±0.07 | -0.08 ±0.12 | 0.56 |
| R398Q + WT | 3.29 ±4.31 |  | 0.67 ±0.07 | 0.02 ±0.12 | 0.47 |
| R398Q 0 mM [Na<sup>+</sup>]<sub>i</sub> | 26.24 ±3.91 | * | 0.91 ±0.10 | 0.58 ±0.13 | -0.09 |
| R928C | -21.84 ±3.47 |  | 0.64 ±0.12 | -0.30 ±0.06 | 0.79 |
| R928C + WT | 11.27 ±6.58 | ** | 0.68 ±0.14 | 0.16 ±0.08 | 0.33 |
| R928C 0 mM [Na<sup>+</sup>]<sub>i</sub> | 26.04 ±4.87 | *** | 0.78 ±0.07 | 0.48 ±0.12 | 0.01 |
| R928C + WT 0 mM [Na<sup>+</sup>]<sub>i</sub> | 41.05 ±6.51 | *** | 0.65 ±0.13 | 0.58 ±0.16 | -0.10 |
| M896I | -23.31 ±5.90 |  | 0.96 ±0.10 | -0.50 ±0.12 | 0.99 |
| M896I + WT | 6.82 ±4.83 | * | 0.65 ±0.06 | 0.12 ±0.06 | 0.37 |
| M896I 0 mM [Na<sup>+</sup>]<sub>i</sub> | 1.78 ±5.58 | * | 0.78 ±0.15 | 0.07 ±0.11 | 0.42 |
| M896I + WT 0 mM [Na<sup>+</sup>]<sub>i</sub> | 43.81 ±6.41 | *** | 0.66 ±0.04 | 0.67 ±0.09 | -0.18 |

**Figure 2:**
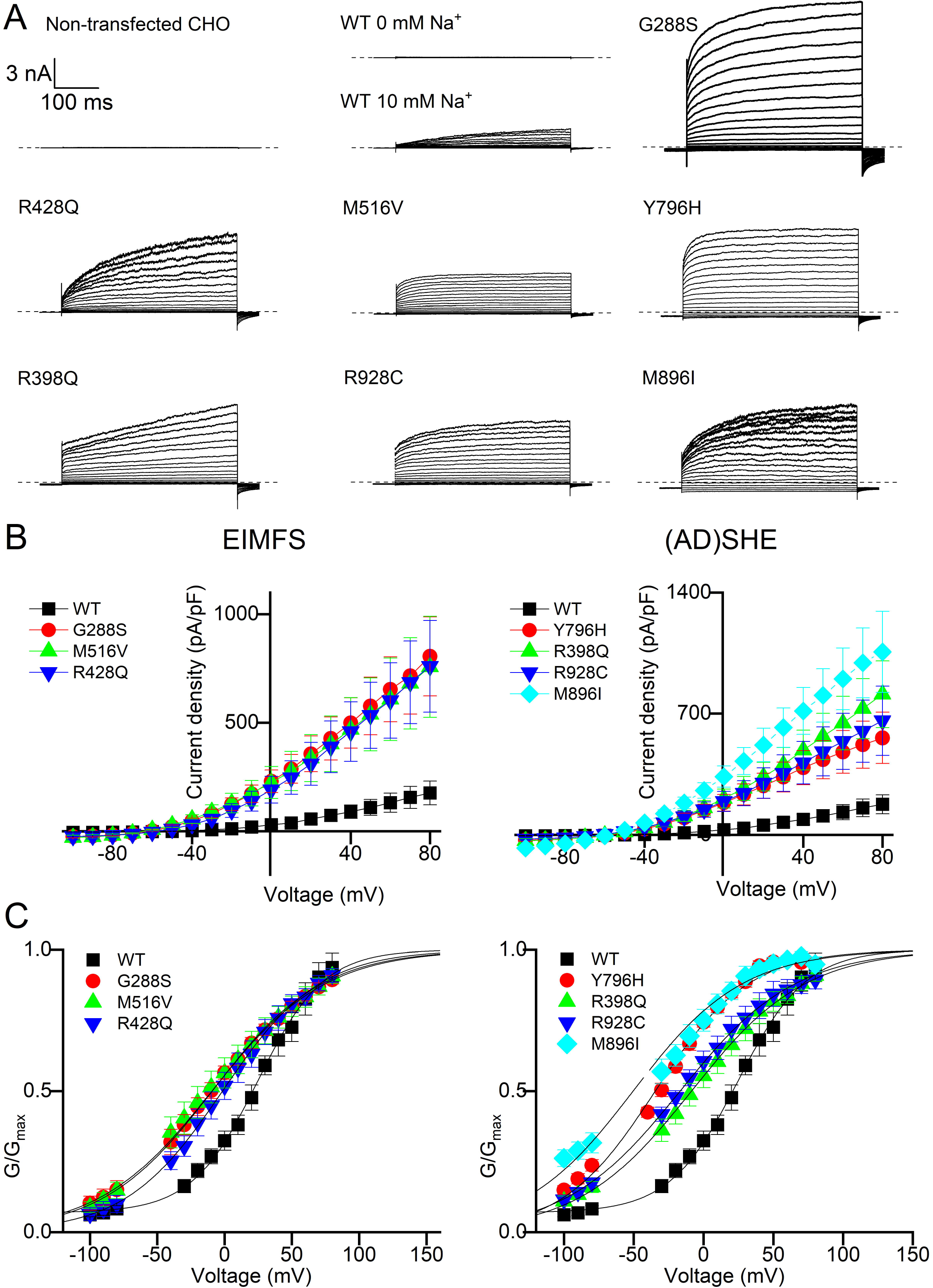
Functional characterisation of mutant K_Na_1.1 carrying EIMFS and (AD)SHE-causing *KCNT1* variants. **A** Representative whole cell currents recorded from non-transfected CHO cells or transfected with wild-type (WT; 0 and 10 mM intracellular Na^+^) or mutant (all 10 mM intracellular Na^+^) K_Na_1.1 as indicated. Currents were recorded in response to 400 ms pulses from −100 to +80 mV in 10 mV increments, from a holding potential of −80 mV. The dashed lines represent the zero-current level. **B** Mean (±SEM) current-voltage relationship for WT K_Na_1.1 (N=6), EIMFS-causing mutant channels (N= 5-9 for each mutant, left panel) and (AD)SHE- causing mutant channels (N= 5-9 for each variant, right panel). **C** Normalised mean (±SEM) conductance-voltage relationship for WT K_Na_1.1 (N=6), EIMFS-causing mutant channels (N= 5-9 for each mutant, left panel) and (AD)SHE- causing mutant channels (N= 5-9 for each mutant, right panel), fitted with Boltzmann functions.

### Co-expressed WT and mutant subunits display a mixture of homomeric WT and mutant characteristics

Co-expression is a method of studying heteromeric assemblies that has been employed in previous functional studies using K_Na_1.1 and closely related K_Na_1.2 ^38,43,44^. The EIMFS-causing mutations; G288S, R428Q and M516V, and (AD)SHE-causing mutations; Y796H, R928C, M896I and R398Q, were co-expressed with WT K_Na_1.1 subunits in a 0.5:0.5 ratio, by plasmid mass, to model the genetic status of heterozygous patients. Mean peak current densities at +10 mV recorded for all mutant+WT co-assemblies with 10 mM intracellular Na^+^ were 3-4-fold larger than homomeric WT channels, regardless of whether the mutation was (AD)SHE or EIMFS-causing (Figure 3A, B and D; Figure 4A, B and D). Only G288S and M896I K_Na_1.1 were significantly smaller than the homomeric mutant when co-expressed with WT K_Na_1.1 subunits (Figure 3D and 4D). Furthermore, with the exception of R398Q+WT, all heteromeric mutant assemblies exhibited a rightward shift in the V_0.5_ value derived from Boltzmann fit of conductance values compared to homomeric mutant channels (Figure 3C, 4C and Table 1).

**Figure 3:**
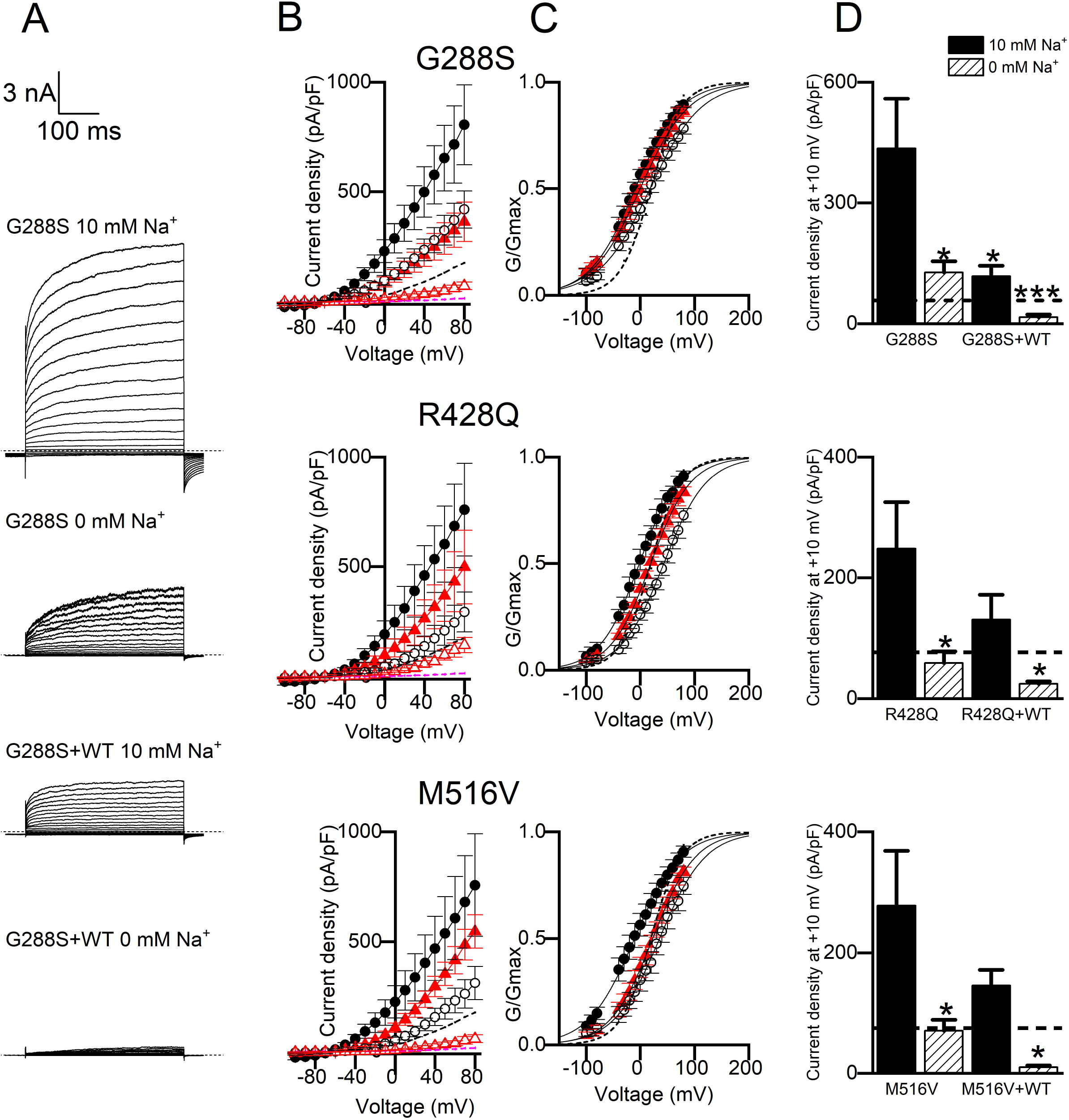
Functional characterisation of EIMFS-causing *KCNT1* variants in the presence and absence of intracellular Na^+^. A Representative currents recorded from homomeric G288S and heteromeric G288S+WT whole cell K_Na_1.1 currents in response to 400 msec steps from −100 to +80 mV in 10 mV increments, from a holding potential of −80 mV in 10 mM and 0 mM intracellular Na^+^, as indicated. Mean (±SEM) current-voltage relationships **(B)** and conductance-voltage relationships fitted with a Boltzmann function **(C)** for EIMFS- causing mutants in 10 mM and 0 mM intracellular Na^+^, and in the presence and absence of co-expressed WT K_Na_1.1 (N=5-9 for all channel types). *Black filled circle*, homomeric mutant with 10 mM Na^+^; *black open circle*, homomeric mutant with 0 mM Na^+^; *red filled triangle*, heteromeric mutant+WT with 10 mM Na^+^; *red open triangle*, heteromeric mutant+WT with 0 mM Na^+^. Mean data for WT K_Na_1.1 with 10 mM Na^+^ indicated with dashed line. **D** Mean peak current amplitude at +10 mV for WT K_Na_1.1 and mutant channels in 10 mM and 0 mM intracellular Na^+^. Mean WT K_Na_1.1 amplitude with 10 mM Na^+^ indicated with dashed line. \**p*<0.05, \*\**p*<0.005, \*\*\**p*<0.0005 compared to homomeric mutant with 10 mM Na^+^, independent one-way ANOVA with Tukey’s post-hoc test (N= 5-9 for each mutant).

**Figure 4:**
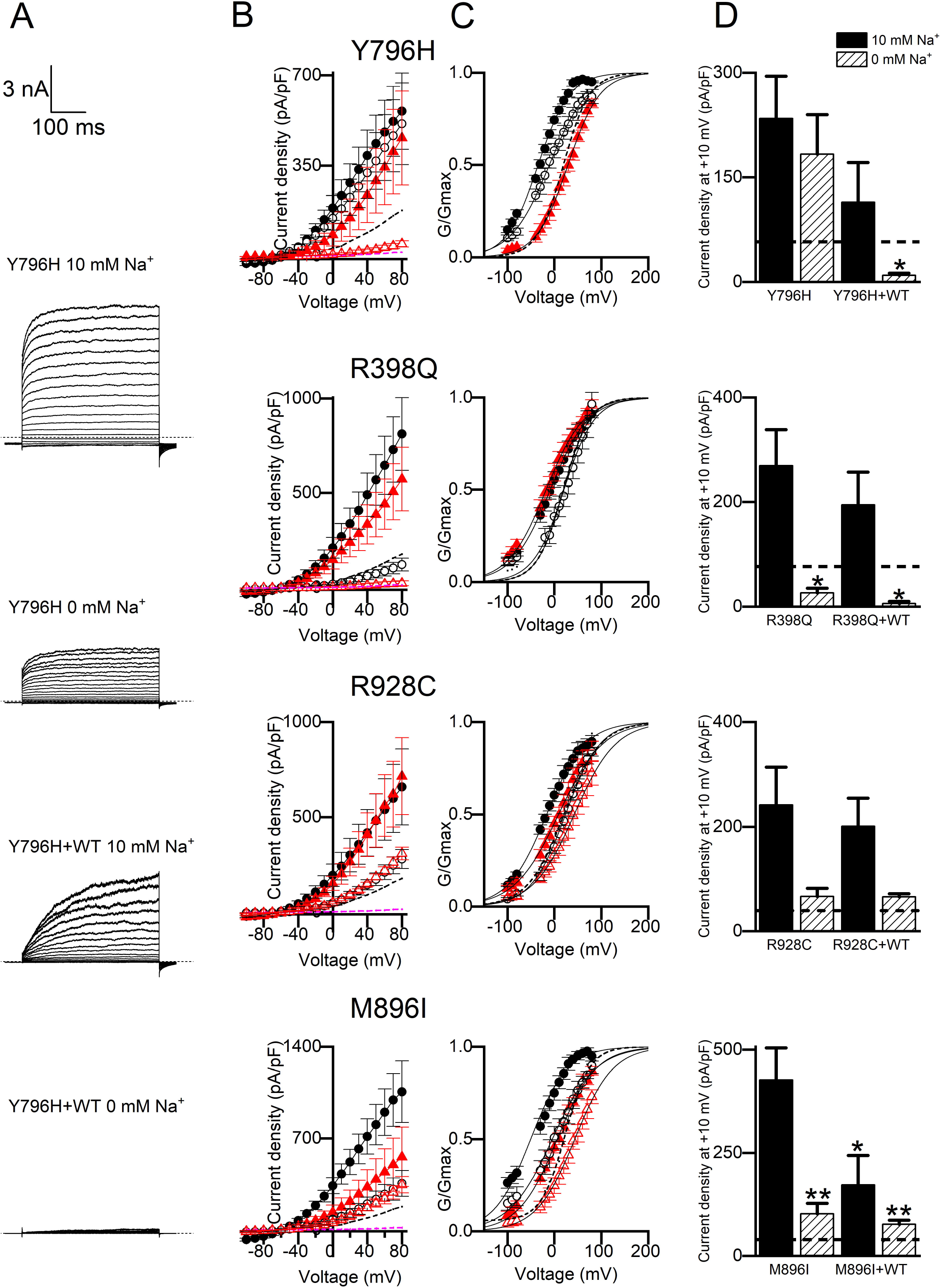
Functional characterisation of (AD)SHE-causing *KCNT1* mutations in the absence of intracellular Na^+^. **A** Representative currents recorded from homomeric Y796H and heteromeric Y796H+WT whole cell K_Na_1.1 currents in response to 400 msec steps from −100 to +80 mV in 10 mV increments, from a holding potential of −80 mV in 10 mM and 0 mM intracellular Na^+^, as indicated. Mean (±SEM) current-voltage relationships **(B)** and conductance-voltage relationships fitted with a Boltzmann function **(C)** for (AD)SHE- causing mutants in 10 mM and 0 mM intracellular Na^+^, and in the presence and absence of co-expressed WT K_Na_1.1 (N=5-9 for all channel types). *Black filled circle*, homomeric mutant with 10 mM Na^+^; *black open circle*, homomeric mutant with 0 mM Na^+^; *red filled triangle*, heteromeric mutant+WT with 10 mM Na^+^; *red open triangle*, heteromeric mutant+WT with 0 mM Na^+^. Mean data for WT K_Na_1.1 with 10 mM Na^+^ indicated with dashed line. **D** Mean peak current amplitude at +10 mV for WT K_Na_1.1 and mutant channels in 10 mM and 0 mM intracellular Na^+^. Mean WT K_Na_1.1 with 10 mM Na^+^ indicated with dashed line. \**p*<0.05, \*\**p*<0.005 compared to homomeric mutant with 10 mM Na^+^, independent one-way ANOVA with Tukey’s post-hoc test (N= 5-9 for each mutant).

### GOF mutant K_Na_1.1 channels activate in a voltage-dependent manner in the absence of intracellular Na^+^

WT K_Na_1.1 channels are primarily activated by intracellular Na^+^, thus removing Na^+^ from the pipette solution would enable any Na^+^-independent activity of mutant K_Na_1.1 channels carrying EIMFS or (AD)SHE-causing *KCNT1* variants to be detected. Na^+^-independent currents have been reported previously for G288S and M516V, and two other EIMFS-causing variants, E893K and R950Q ^38,43^. No (AD)SHE-causing variants or other DEE-causing variants, including R428Q, have been studied in this way. Each of the homomeric mutant channels examined activated in the absence of Na^+^, albeit with smaller current densities than those recorded in the presence of intracellular Na^+^. The decrease in mean peak current density at +10 mV was modest, with a 1-4-fold decrease for all mutants with the exception of R398Q, which showed a 13-fold decrease (Figure 3D and 4D). Notably, when conductance values derived from the recorded current amplitudes in the absence of Na^+^ were fitted with a Boltzmann equation (Equation 2), these were shifted by 15-60 mV in the depolarising direction compared to activation curves obtained with 10 mM Na^+^ (Figure 3B, 4B and Table 1). For all mutants except for Y796H, the activation midpoint became positive in the absence of Na^+^ (Table 1).

For co-expressed mutant+WT K_Na_1.1 subunits, there was a resulting 7-34-fold decrease in the mean peak current density recorded at +10 mV for all mutant+WT channels in the absence of intracellular Na^+^ compared to currents recorded with 10 mM intracellular Na^+^ (Figure 3D and 4D). Although small K_Na_1.1 currents were recorded, the size of the currents prevented Boltzmann analysis of most mutant+WT channels in the absence of intracellular Na^+^. Two (AD)SHE-causing mutants, M896I and R928C, produced sufficiently large currents in the absence of Na^+^ when co-expressed with WT K_Na_1.1. The V_0.5_ value in both cases was more positive than the homomeric mutant under similar conditions (Figure 4C and Table 1).

### The time constant of channel activation of both WT and mutant K_Na_1.1 is independent of voltage and Na^+^

To examine additional effects that *KCNT1 GOF* mutations may have on K_Na_1.1 activation kinetics, single exponential functions were fitted to the currents at each voltage measured to yield the activation time constant (*τ*, from Equation 3) (Figure 5A). Notably, although the channels have weak voltage sensitivity and time-dependent activation, the time constants obtained from both WT and mutant channels were voltage-independent, with no significant difference across the voltage range (Figure 5B). The K_Na_1.1 activation time constant of was not significantly altered by most mutations studied. An exception was (AD)SHE-causing K_Na_1.1 mutation Y796H, with which the time constant decreased 2-fold at +10 mV (Figure 5B and C). Another (AD)SHE-causing mutation, R398Q, could not be reliably fit with a single or double exponential function, but appeared to activate considerably more slowly than WT K_Na_1.1 channels. This is consistent with characterisation of channels expressed in *Xenopus* oocytes showing this mutant to have a slower time-to-peak ^41^. R928C (Figure 5A) and M896I currents could not be fit adequately with a single exponential function, and were better fit with two exponentials, giving a fast (**τ**_fast_) and slow (**τ**_slow_) time constant (Figure 5C).

**Figure 5:**
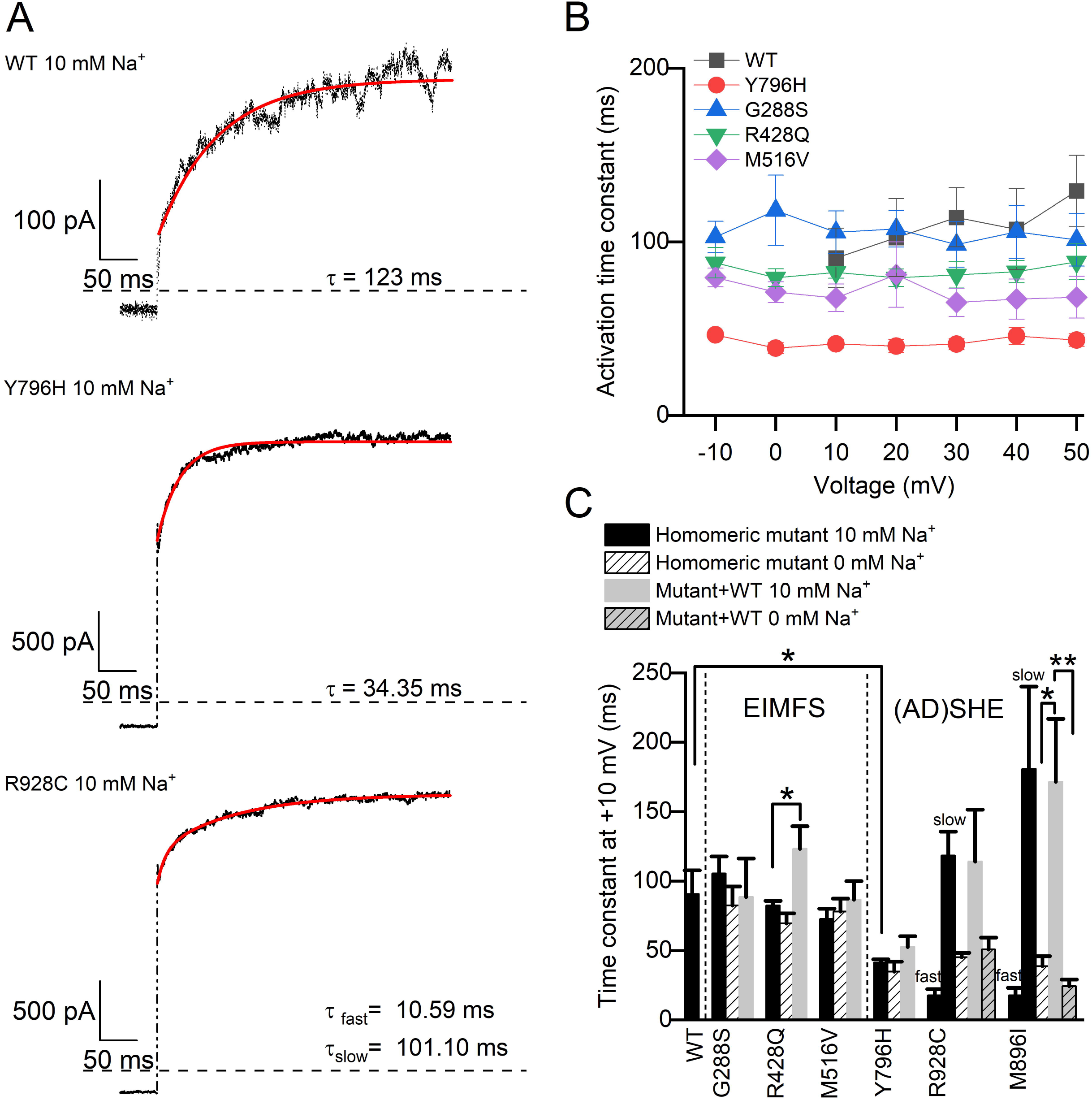
Activation time-course of WT and mutant K_Na_1.1 in the presence and absence of intracellular Na^+^. **A** Representative currents recorded from WT and Y796H K_Na_1.1 channels (black) fitted with a single exponential function (red), and R928C K_Na_1.1 with a bi-exponential function (red), recorded at +10 mV with 10 mM intracellular Na^+^. **B** Summary of mean (±SEM) time constant (τ) values derived from a single exponential fit of homomeric WT K_Na_1.1, (AD)SHE-causing and EIMFS-causing mutants obtained at different voltages in 10 mM intracellular Na^+^ (N= 5-9 for each mutant). Only those mutants that yielded mono-exponential time constants are included. **C** Mean (±SEM) τ values at +10 mV for EIMFS- and (AD)SHE-causing homomeric mutant and co-expressed heterometic mutant K_Na_1.1 channels in 10 mM and 0 mM intracellular Na^+^. WT K_Na_1.1 is shown only with 10 mM Na+ and mutant K_Na_1.1 data are grouped by mutation as indicated. Where R928C and M896I currents were fit with a bi-exponential function, τ fast and τ slow are indicated above the bars. \**p*<0.05, \*\**p*<0.005 independent one-way ANOVA with Tukey’s post-hoc test compared to homomeric mutant with 10 mM Na^+^ (N= 5-9 for each mutant).

Currents recorded from CHO cells expressing DEE-causing mutant K_Na_1.1 in the absence of intracellular Na^+^ were also fit with a single exponential function. As with 10 mM Na^+^ in the pipette solution, the **τ** values were similar at all voltages measured. The **τ** values were unchanged from those in the presence of Na^+^, implying that removal of Na^+^ from the intracellular solution does not affect the slowly activating, time-dependent component of channel activation (Figure 5C). The only exceptions to this were R928C and M896I (AD)SHE-causing mutations, which were previously fit best with two exponential components. In the absence of Na^+^ however, the currents could be adequately fit with a single exponential function (Figure 5C).

All currents produced by co-expression of WT and mutant subunits were adequately fit with a single exponential, except for R398Q+WT, which could not be fit as a homomeric R398Q channel previously. Aside from Y796H+WT, similar to homomeric mutant channels, the **τ** values obtained for all mutant+WT heteromers were unchanged from WT across all voltages tested (Figure 5C). In the heteromeric assembly, both M896I and R928C were again better fit with a single exponential in both the absence and presence of intracellular Na^+^. With 10 mM Na^+^ the **τ** value was reflective of the WT channel, and in the absence of Na^+^ it was faster and more reflective of the homomeric mutant (Figure 5C).

### Selectivity filter gating, and not Na^+^-dependence, is altered by mutation of T314

The voltage- and Na^+^-independent activation time constants were reminiscent of K^+^ channel subtypes that exhibit voltage-dependent selectivity filter gating, such as several mammalian tandem pore domain K^+^ channels (K2P) ^32,33,53–55^. In outwardly-rectifying K2P channels, mutation of a threonine at the cytoplasmic end of the selectivity filter motif that forms the intracellular-most K^+^ binding site resulted in voltage- and ligand-independent channel currents ^32^. The equivalent threonine residue in the K_Na_1.1 selectivity filter motif is T314, which we mutated to cysteine (T314C). This mutation also resulted in voltage-independent currents, but these reversed close to 0 mV (Fig. 6). Importantly, the Na^+^-sensitivity of T314C K_Na_1.1 remained intact since, like WT K_Na_1.1, currents were recorded in the presence of 10 mM intracellular Na^+^ but were negligible in the absence of intracellular Na^+^ (Fig. 6). These effects of mutating the putative selectivity filter gate were distinct from the mutations caused by the DEE-causing *KCNT1* GOF variants.

**Figure 6:**
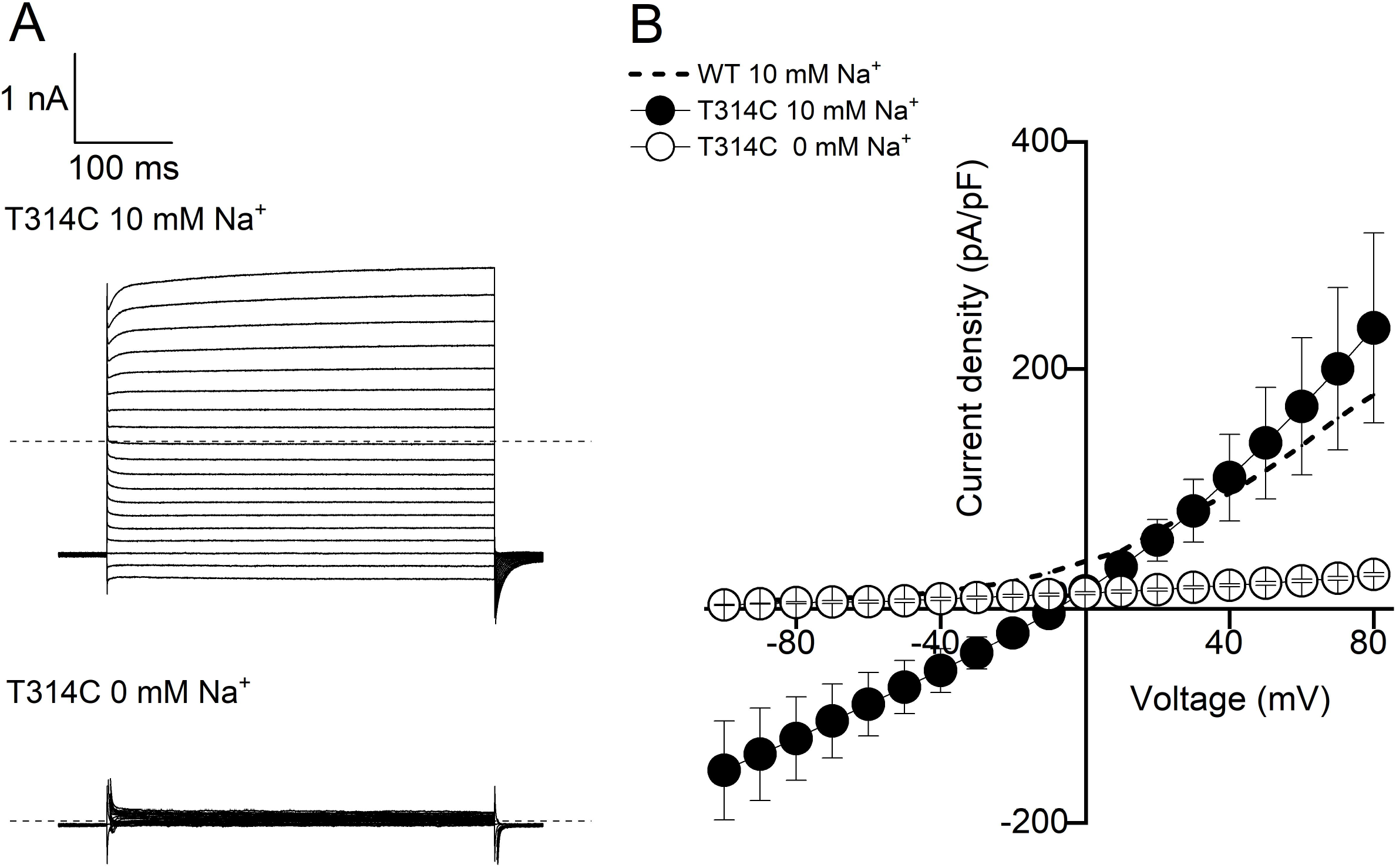
Mutation of a conserved threonine residue in the K_Na_1.1 selectivity filter disrupts selectivity and voltage-activation, but not Na^+^-activation. **A** Representative whole cell currents recorded from T314C K_Na_1.1 in response to 400 ms steps from −100 to +80 mV in 10 mV increments, from a holding potential of −80 mV in 10 mM and 0 mM intracellular Na^+^ as indicated. **B** Mean (±SEM, N=5) current-voltage relationships for T314C K_Na_1.1 in 10 mM and 0 mM intracellular Na^+^. Data for mean WT K_Na_1.1 current-voltage relationship indicated with dashed line.

## Discussion

Missense *KCNT1* variants result in amino acid substitutions in several regions of the K_Na_1.1 protein structure. All these DEE-causing mutations result in increased channel activity and so we hypothesised that this, at least in part, is mediated by a common mechanism. To address this, we studied the activation properties of a range of missense mutations that cause different severities of human disease and result in amino acid substitutions in different regions of the channel. We found that unlike WT K_Na_1.1, the channels formed from mutant subunits could each be activated in a voltage-dependent manner in the absence of intracellular Na^+^. We propose that this a feature common to all gain-of-function K_Na_1.1 mutations caused by pathogenic variants. On the mechanism by which missense mutations in diverse regions of the channel protein converge to bring about Na^+^-independent gating, the closely related K_Ca_1.1 (BK_Ca_, KCNMA1) channel may provide some clues. It was previously proposed that the regions in the RCK domains of the K_Ca_1.1 channel C-terminal act as a negative modulator of channel opening, and undergo a conformational change following Ca^2+^ binding to relieve inhibition ^56^. This form of autoinhibition may similarly exist in K_Na_1.1, but with the conformational changes associated with Na^+^-binding ^28^ relieving inhibition and allowing the activation gate to open. Then this would explain how mutations affecting different parts of the channel could cause Na^+^-independent channel opening, despite many being located distally from the pore-forming region. It is possible that these mutations destabilise the inactive unliganded channel conformation, making it easier for the activation gate to open by lowering the activation energetic barrier. These disease-related mutations could therefore be considered as loss-of-function with respect to this autoinhibitory or inactivation mechanism. A schematic of this gating process is illustrated in Figure 7.

**Figure 7:**
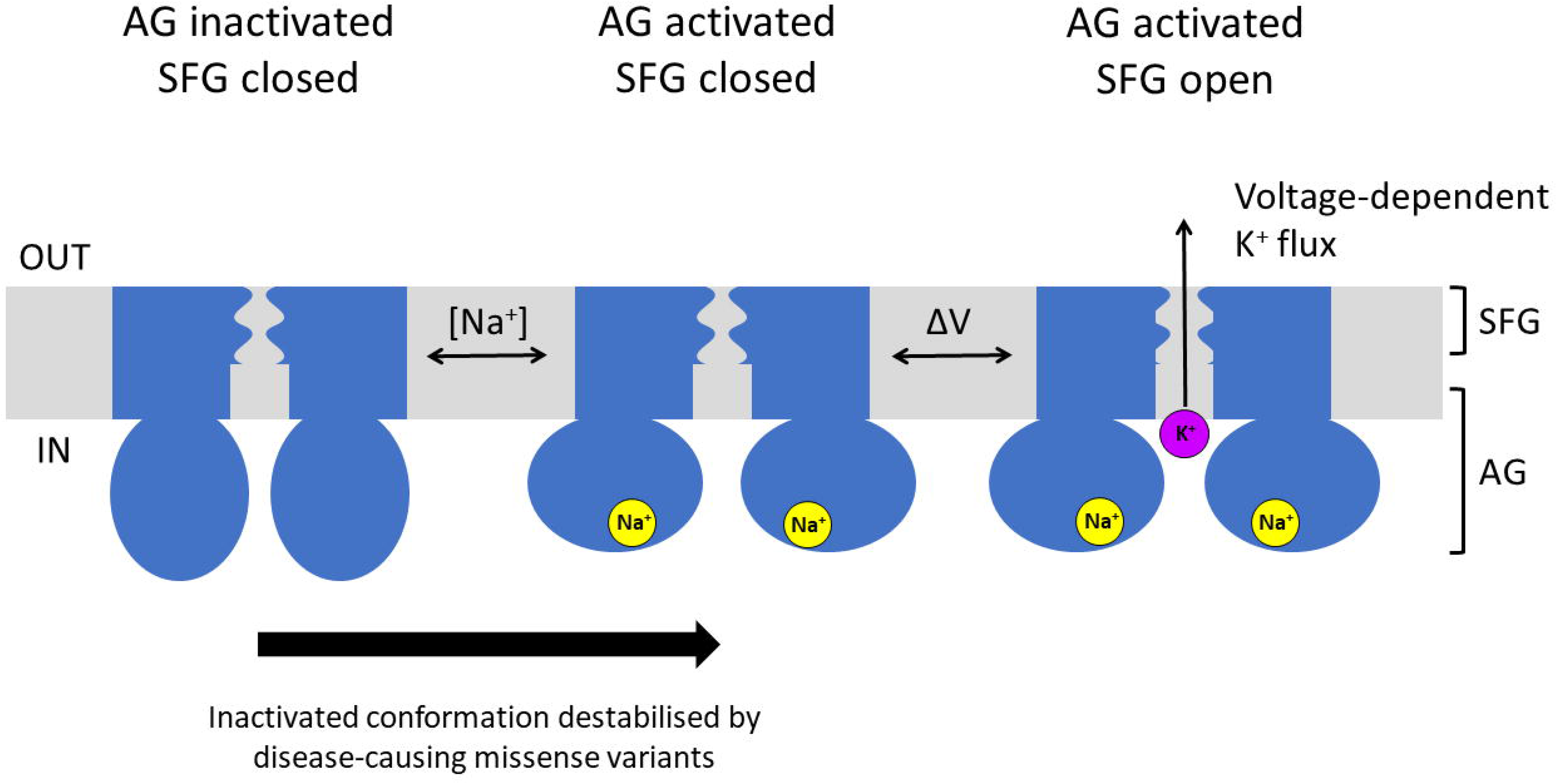
Schematic of K_Na_1.1 gating and effects of disease-causing mutations. The K_Na_1.1 channel gating is governed by an activation gate (AG) that is modulated by intracellular Na^+^ and a selectivity filter gate (SFG) that is opened by voltage-dependent K^+^ flux once the channel is in the activated conformation. The primary effect of disease-causing variants is hypothesised to destabilise the inactivated channel conformation (*left*). This lowers both the free energy and, indirectly, the dependence on Na^+^ for the channel to become activated (*middle*) and permit voltage-dependent K^+^ currents (*right*).

Conceptionally, the idea of a Na^+^-dependent activation gate, similar to the activation gate of MthK and other K^+^ channels ^33^, provides a mechanistic understanding of how mutations increase K_Na_1.1 channel activity. Na^+^ binding to the intracellular K_Na_1.1 domains and/or the presence of GOF-causing amino acid substitutions lower the energetic barrier between the inactive and active channel states. We quantified this using the zero-voltage free energy calculation for channel opening (Table 1). As there were no detectable WT K_Na_1.1 currents in the absence of intracellular Na^+^, this means that the activation energy barrier for unliganded channels is high, and is then lowered in a Na^+^-dependent manner. The voltage-dependent activation measured for each of the mutant K_Na_1.1 channels in the absence of intracellular Na^+^ indicates that the activation energy barrier of unliganded channels is relatively low. This energy barrier is lowered further in an additive manner by intracellular Na^+^, as indicated by the shift of V_0.5_ to more negative potentials and the more negative ΔG_0_ with 10 mM compared to 0 mM Na^+^ (Table 1). Co-expression of WT and mutant K_Na_1.1 yielded currents with properties intermediate of WT and mutant subunits alone, suggesting that each subunit in the tetramer contributes to channel activation, possibly in an additive manner. This mechanism for increased channel activity does not require a change in the Na^+^-binding affinity or mechanism for Na^+^-activation, but would give the appearance of increased Na^+^ sensitivity and a shift in EC_50_ to lower concentrations as seen by Tang *et al* ^36^. Lowering of the activation barrier by mutations also explains the increase in co-operative channel opening, compared to WT channels, observed in excised patches containing multiple K_Na_1.1 channels ^3^.

K^+^ channels that lack a gating mechanism though an S6 helix bundle-crossing usually feature a gate formed by the selectivity filter ^32,33,53,54,57^. It has been proposed that the structure of the selectivity filter changes as a result of allosteric coupling with the activation gate, facilitated by pore-lining transmembrane helices, to facilitate ion conduction ^33^. Importantly, a threonine residue at the cytoplasmic end of the selectivity filter is thought to be critical in the gating of MthK, K2P, K_Ca_1.1 and KcsA channels ^32,33,53,57^. This threonine residue is conserved in K_Na_1.1 and the T314C mutation disrupted rectification. We found that this mutation affected permeation and voltage-sensitivity but retained its Na^+^-sensitivity and yielded negligible current in the absence of intracellular Na^+^, similar to WT K_Na_1.1. This reinforces the idea that the activation gate, and not the selectivity filter gate, is directly modulated by Na^+^-binding to the channel protein, and it is the activation gate that is predominantly affected by DEE-causing GOF mutations. It is therefore likely that the Na^+^ and voltage-independent time constant for K_Na_1.1 channel activation represents the time course of the events in the selectivity filter that follow Na^+^ and voltage-activation, resulting in ion conduction. Voltage-independent activation time constants have also been observed with the activation of K2P channels that exhibit voltage dependence and selectivity filter gating ^32^. It is further possible that some disease-causing *KCNT1* mutations also affect selectivity filter gating, in addition to their effects on the activation gate. (AD)SHE-related Y796H K_Na_1.1 activated over a faster time course (reduced τ) than WT K_Na_1.1. Increased maximum open probability (P_O_) has previously been found with a number of K_Na_1.1 mutants, including Y796H ^36^ and it is possible that faster activation time constants correlate with increased maximum P_O_. For transitions between an open and closed state, a faster time constant arises from increase in either the opening or closing rate constant, however an increase in maximum P_O_ is consistent only with an increase in the opening rate constant. Consistent with this, (AD)SHE-causing variant R398Q was reported as having a decreased P_O_ at 25 and 50 mM intracellular Na^+^ ^36^, and we found this variant to activate very slowly, preventing reliable fitting with either a mono- or bi-exponential function. Milligan *et al* previously described this variant as having a slower time-to-peak in *Xenopus* oocytes ^41^. Exceptions to the mono-exponential activation were found with R928C and M896I K_Na_1.1 where a second and faster exponential was detected in the presence of intracellular Na^+^. These mutations may slow a relatively fast Na^+^-dependent activation of the channel, revealing this otherwise undetectable step. It is unclear however, the degree to which these additional changes to channel kinetics contribute to the GOF.

To conclude, we have found that each of the DEE-causing *KCNT1* variants studied here lower the energetic requirements for ligand-dependent activation, which manifest as K_Na_1.1 channel activation in the absence of intracellular Na^+^ and overall GOF. We propose that each of the amino acid substitutions across the channel structure that result in channel GOF destabilise the inactive unliganded channel conformation. Through co-expression with WT subunits, we have demonstrated that heteromeric channels do not behave the same as homomeric mutant channels but have less-pronounced shifts in activation kinetics and activity in the absence of intracellular Na^+^. Previous work characterising homomeric mutant channels, therefore, should be interpreted with caution when postulating how endogenous channels containing mutant K_Na_1.1 subunits behave in vivo, how this affects neuronal excitability, and how this may lead to epilepsy phenotypes. Our results suggest that heteromeric channels differ in their kinetic behaviour to homomeric mutations and thus will have distinct effects on neuronal excitability. Previous work has found quinidine to be a more potent inhibitor of some mutant channels compared to WT K_Na_1.1, possibly due to an open channel block mechanism, and which may not be as pronounced in heteromeric assemblies of mutant and WT subunits ^38,41,43,51^. Whilst the method of studying the heteromeric assembly has limitations, it may provide a more physiologically relevant model for studying the functional implications of DEE-causing *KCNT1* variants on K_Na_1.1 channel behaviour. This will also be advantageous for development of pharmacological interventions, when studying the inhibition of either homomeric WT or mutant channels *in vitro*, where any differences in observed inhibitor potency may be of limited clinical relevance.

## Abbreviations

(AD)SHE: autosomal-dominant or sporadic sleep-related hypermotor epilepsy
DEE: developmental and epileptic encephalopathy
EIMFS: epilepsy of infancy with migrating focal seizures
GOF: gain-of-function
K2P: two pore domain-containing potassium channel subunit
K_Na_1.1: sodium-activated potassium channel subunit
P_O_: channel open probability
WT: wild-type

## Acknowledgments

Supported by a BBSRC-CASE PhD studentship awarded to B.A.C. (BB/M011151/1) and Autifony Therapeutics Ltd.

## Notes

<u>Conflicts of Interest statement</u> The authors have no conflicts of interest to declare.

### Competing Interest Statement

The authors have declared no competing interest.

### Summary of Updates

Main text edited for typos, font errors, and clarity. Summary scheme added as Figure 7.

